# An ecological investigation of the capacity to follow simultaneous speech and preferential detection of ones’ own name

**DOI:** 10.1101/2022.06.07.495173

**Authors:** Danna Pinto, Maya Kaufman, Adi Brown, Elana Zion Golumbic

**Affiliations:** The Gonda Multidisciplinary Center for Brain Research Bar Ilan University, Israel

**Keywords:** Divided Attention, Speech Processing, Cocktail Party, Dual-Task

## Abstract

Many situations require focusing attention on one speaker, while monitoring the environment for potentially important information. Some have proposed that dividing attention among two speakers involves behavioral tradeoffs, due to limited cognitive resources. However the severity of these tradeoffs, particularly under ecologically-valid circumstances, is not well understood. We investigated the capacity to process simultaneous speech using a dual-task paradigm simulating task demands and stimuli encountered in real-life. Participants listened to conversational narratives (Narrative Stream) and monitored a stream of announcements (Barista Stream), to detect when their order was called. We measured participants’ performance, neural activity and skin conductance as they engaged in this dual-task.

Participants achieved extremely high dual-task accuracy, with no apparent behavioral tradeoffs. Moreover, robust neural and physiological responses were observed for target-stimuli in the Barista Stream, alongside significant neural speech-tracking of the Narrative Stream. These results suggest that humans have substantial capacity to process simultaneous speech and do not suffer from insufficient processing resources, at least for this highly ecological task-combination and level of perceptual load. Results also confirmed the ecological validity of the advantage for detecting ones’ own name at the behavioral, neural and physiological level, highlighting the contribution of personal relevance when processing simultaneous speech.

## Introduction

The brain’s capacity and bottlenecks for processing the speech of multiple concurrent talkers has been highly debated by neuroscientists and cognitive psychologists. Research into this question has largely focused on selective attention paradigms - where participants are instructed to listen to one ‘main’ stimulus and perform a task, while disregarding other task-irrelevant stimuli (Cherry, 1953; Broadbent, 1958; Lane and Pearson, 1982; Driver, 2001; Ding et al., 2018). However, these selective attention paradigms have yielded mixed results regarding the depth of processing that is applied to task-irrelevant speech, with some studies suggesting that task-irrelevant speech is only represented at an acoustic level but not at a semantic/linguistic level, while others do find evidence for some linguistic processing of task-irrelevant speech (Dupoux et al., 2003; Brodbeck et al., 2020; Dai et al., 2021; Har-shai Yahav and Zion Golumbic, 2021), particularly if it contains salient content words (Moray, 1959; Treisman, 1960; Wood and Cowan, 1995; Rivenez et al., 2008) Perhaps one of the most well-known demonstrations of this phenomena is the conscious detection of one’s own name in supposedly “unattended” speech (Moray, 1959; Wood and Cowan, 1995; Conway et al., 2001; Tamura et al., 2012; Tateuchi et al., 2012; Röer et al., 2013; Naveh-Benjamin et al., 2014; Holtze et al., 2021). However, one major tension in interpreting these results is the ambiguity regarding what participants actually do in selective attention tasks. Some have suggested that the reported effects of detecting words in task-irrelevant speech result from momentary shifts of attention between the streams (glimpsing), rather than true processing of unattended speech (Holender, 1986; Lachter et al., 2004; Brungart and Iyer, 2012). This problem is exacerbated by the fact that in selective attention designs the behavioral readouts regarding the task-irrelevant stimulus are indirect at best (Driver, 2001; Tun et al., 2002; Dupoux et al., 2003; Beaman, 2004; Humes et al., 2006; Rivenez et al., 2006; Carey et al., 2014; Aydelott et al., 2015; Schepman et al., 2016). This ‘ground-truth’ uncertainty is inherent to all selective attention paradigms, making it difficult to disentangle the interaction between participants’ internal state of attention and the depth of speech processing.

Besides the inherent ‘ground-truth’ ambiguity of selective attention paradigms, it is not clear to what extent it captures the type of listening prioritization/strategy that is naturally employed in multi-talker contexts. Arguably, in many real-life contexts it can be beneficial for listeners to monitor secondary speech streams for potentially relevant information, rather than attempting to suppress or ignore them (Kahneman and Treisman, 1984; Näätänen, 1988). Therefore, here we ask to what extent is the system capable of processing two concurrent speech streams when they are both task-relevant. To this end, we designed a dual-task ‘cocktail-party-style’ paradigm that resembles the type of listening challenges encountered in real-life multi-speaker contexts. Our design simulates the experience of being in a café and listening to a narrative of conversational speech from your partner (Narrative Stream), while <u>also</u> monitoring a background stream of café-order announcements (e.g. “coffee for David; Barista Stream) and responding when your order is called. This design not only circumvents the need to make assumptions about how attention is allocated between the two speakers, but provides explicit behavioural measures about both stimuli, allowing us to assess potential ‘trade-offs’ between them.

Relative to previous studies of distributed attention to speech, where listeners were asked to listen to two concurrent speech streams and perform the same task on both (e.g. detect targets, Lambez et al., 2020; Agmon et al., 2021, Brungart et al., 2001, 2005; Best et al., 2005, 2006, 2010a; Shafiro and Gygi, 2007a; Abel et al., 2012; Gygi and Shafiro, 2014a; Koelewijn et al., 2014; Baldock et al., 2019 or answer comprehension questions, Boudewyn and Carter, 2018; Kaufman and Zion Golumbic, 2022), the current design recognizes that concurrent speech may be relevant to the listener in different ways. The specific task-combination used here – listening to the Narrative Stream for its content and monitoring the Barista Stream for target-words – implicitly designate the streams as primary and secondary, respectively. The realistic nature of these task-demands, together with the highly contextual speech materials used here, emulate the type of listening demands that listeners may encounter in real-life multi-speaker contexts, enhancing the ecological validity of this study.

This unique design also afforded us the opportunity to revisit another key research question, regarding the unique-status of hearing one’s own name in multi-speaker contexts (the “Cocktail Party Effect”; (Cherry, 1953; Moray, 1959; Wood and Cowan, 1995; Conway et al., 2001; Tamura et al., 2012; Tateuchi et al., 2012; Röer et al., 2013; Naveh-Benjamin et al., 2014; Holtze et al., 2021). There is ample evidence in the literature of unique behavioral, physiological and neural responses to hearing ones’ own name vs. other words or names, when these are presented alone (Berlad and Pratt, 1995; Folmer and Yingling, 1997; Müller and Kutas, 1997; Perrin et al., 2006, 1999, 2005; Höller et al., 2011; Del Giudice et al., 2014; Lechinger et al., 2016; Liu et al., 2019; Jijomon and Vinod, 2021). However, the robustness of this own name advantage in cocktail-party-style experiments, and particularly when the name appears in a task-irrelevant stream, has been more difficult to assess and replicate, and varies substantially across individuals as a function of age and working-memory capacity (Conway et al., 2001; Eichenlaub et al., 2012; Tamura et al., 2012; Tateuchi et al., 2012; Naveh-Benjamin et al., 2014; Holtze et al., 2021). The dual-task design used here lends itself nicely to revisiting this effect under highly ecological conditions, by sometimes using the participants’ own name as the target name in the Barista Stream and testing whether responses were enhanced relative to non-personally relevant names.

Participants’ neural activity (electroencephalography; EEG) and physiological responses (galvanic skin response; GSR) were recorded throughout the experiment. Behavioral performance on the dual-task and analysis of the neural speech-tracking response to the Narrative and Barista Streams were used to assess the nature of potential ‘trade-offs’ in processing these concurrent speech stimuli which were both task-relevant, but in different ways (Kerlin et al., 2010; Ding and Simon, 2012a; Mesgarani and Chang, 2012; Power et al., 2012; Zion Golumbic et al., 2013; O’Sullivan et al., 2015; Fuglsang et al., 2017; Fiedler et al., 2019; Kaufman and Golumbic, 2022). We also studied the time-locked neural and physiological responses to target-words in the Barista Stream, complementing the behavioral data and providing insight into the potential ‘special status’ of detecting ones’ own name in a secondary speech stream. This combined behavioral, neural, and physiological data set, coupled with the use of a highly ecological task and stimuli, provides a broad perspective on the cognitive and neural mechanisms involved in the attempt to actively follow two speech streams at the same time.

## Methods

### Participants

41 participants took part in this study; however, 5 participants were excluded due to technical issues. Therefore, the reported analyses include a total of 36 participants (22 female; 14 male; ages 19-30, mean age 24). All participants were fluent Hebrew speakers, with self-reported normal hearing and no history of psychiatric or neurological disorders. Participants were paid or received course credit for participation. The study was approved by the Ethics Committee of Bar-Ilan University, and all participants read and signed an informed consent form prior to the commencement of the experiment.

### EEG Recording and Apparatus

EEG was recorded using a 64 Active-Two system (BioSemi) with Ag-AgCl electrodes, placed according to the 10-20 system, at a sampling rate of 1024 Hz. Additional external electrodes were used to record from the mastoids bilaterally and both vertical and horizontal EOG electrodes were used to monitor eye-movements. The experiment was conducted in a dimly lit acoustically and electrically shielded booth. Participants were seated on a comfortable chair and were instructed to keep as still as possible and breathe and blink naturally. Experiments were programmed and presented to participants using PsychoPy (https://www.psychopy.org) (Peirce et al., 2019). Visual instructions were presented on a computer monitor, and auditory stimuli were delivered through in-ear earphones (Etymotic ER-1). Button-press responses were recorded using a keyboard

### GSR Recording

Skin conductance or Galvanic skin response (GSR) was measured using two passive Nihon Kohden electrodes placed on the fingertips of the index and middle fingers of participants’ non-dominant hand. The signal was recorded through the BioSemi system amplifier and was synchronized to the sampling rate of the EEG.

### Speech Materials

Two types of speech materials were presented in this study, which we refer to as the ‘Narrative Speech’ and ‘Barista Speech’. Material for both speech streams were recorded in-house in a sound attenuated booth.

#### Narrative speech

Materials for the Narrative Stream were recorded by a male actor. To encourage spontaneous conversational-style speech, the actor was given a series of prompts and was asked to speak about them for approximately 40 seconds (timer shown on screen). The prompts referred primarily to daily experiences, and could be either of a personal nature (e.g. “describe a happy childhood memory”) or more informational (e.g. “describe how to prepare your favorite meal”). The actor was given the list of prompts in advance in order to plan their answers, however the narratives themselves were delivered in a spontaneous, unscripted fashion. The original recordings were edited slightly to ensure continuous speech, by removing filler words (e.g., ‘umm’, ‘ugh’) and shortening long pauses. The amplitude of all narrative recordings was equated using the software PRAAT (Boersma and Weenink, 2010). In total, we used 40 narrative recordings, ranging between 31-37 seconds in length.

#### Barista Speech

The Barista Speech consisted of lists of orders and filler sentences that might be heard in café. The stimuli were constructed from single words, recorded individually by a female actor, in no systematic order. Individual words were cut and equated for amplitude (using PRAAT), and then concatenated (using Matlab) to create order-sentences, that consist of a person’s name and a food-item they had ordered. Order-sentences had two possible syntactic structures, which in English correspond to “Salad for Sara”, or “Sara’s salad is ready”. Importantly, in Hebrew, in both these structures the name appears in the 3^rd^ word position of the sentence (“Salat bishvil Sara” vs. “ha-salat shel Sara muckan”), and hence its occurrence is similarly predictable. Besides order-sentences, additional filler sentences were constructed, that did not contain any names or food items, but are contextually relevant for a café environment (e.g., “Clean-up needed at table five”, “Lunch served until four”). In all sentences, words were presented with a constant between-word interval of 100ms. Individual sentences were further concatenated into streams (Figure 1C), with a 200ms between-sentence interval to create streams that matched the narratives in length (each containing 10-13 sentences).

**Figure 1:**
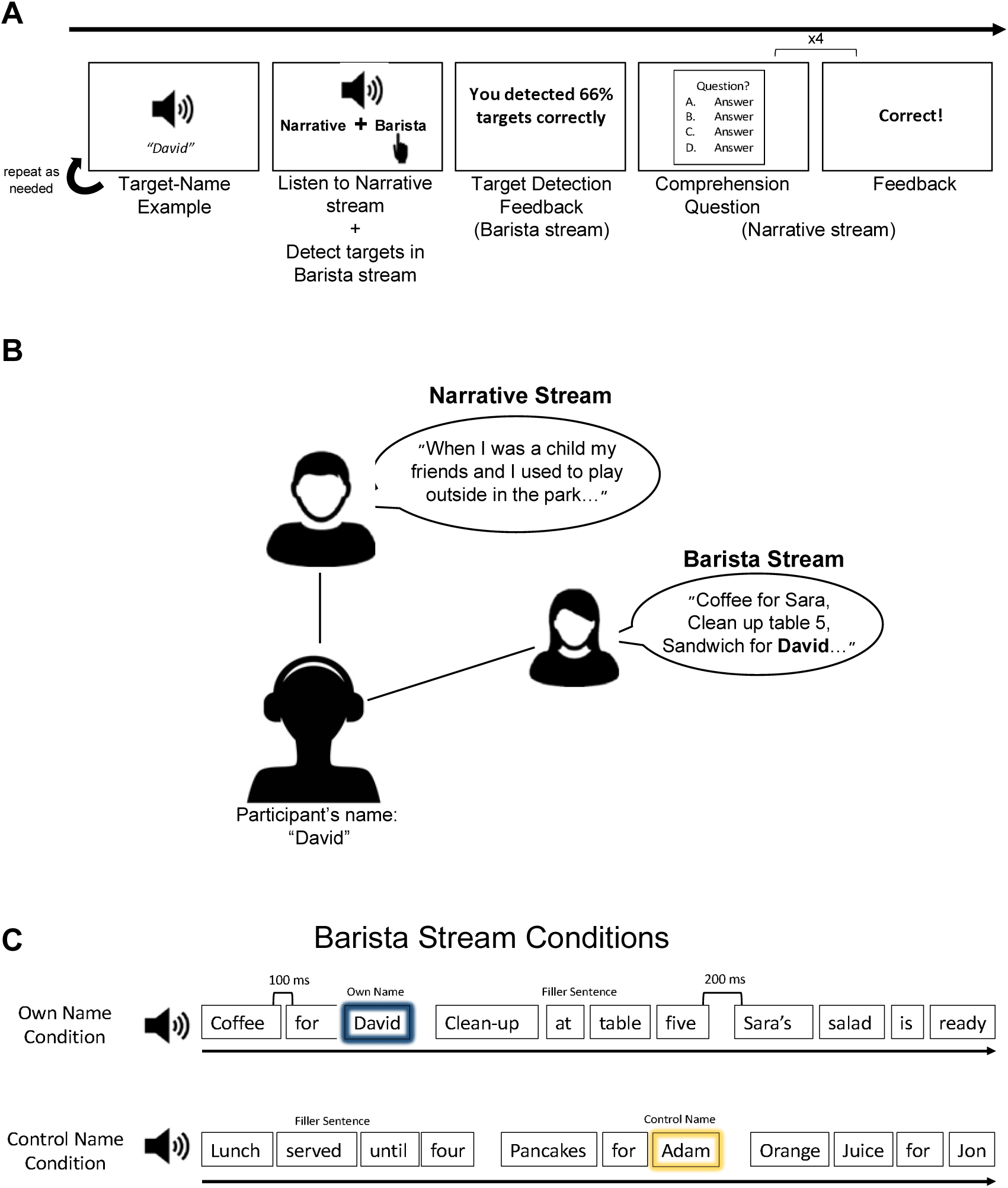
Illustration of the dual-task paradigm. A. The time course of individual trials. At the start of the trial, participants were familiarized with the audio recording of the target name in that trial (their own / control name). Then, the two streams were presented concurrently (as shown in B), and participants were instructed to listen to the content of the Narrative Stream and also respond via button press when they heard the target name in the Barista Stream. After hearing the speech, participants received feedback regarding their target detection accuracy and proceeded to answer four multiple-choice questions regarding the content of the Narrative Stream. B. Illustration of the dual-task design. The Narrative Stream, spoken in a male voice, was presented from a central location whereas the Barista Stream, in a female voice, was presented at a reduced intensity and only to the right ear. C. Illustration of the stimuli used for the Barista Stream, in the own name and control conditions. The Barista Stream consisted of order-sentences (containing names and food items) and filler sentence. The target name was either the participants’ own name or their personal control name.

The sentences and streams were personalized for each participant, in order to investigate the own name advantage. For each participant two names were selected as targets – their name (’own name’) and the name of the previous participant in the experiment (‘control name’). This ensured that, across participants, acoustic differences between names were orthogonal to their status as own/control. For each participant, 40 personalized streams were constructed, of them - half included order-sentences with their own name and half with their control name, serving as target words. Target words could be presented between 1-3 times in a given stream, but the own name and control name targets never co-occurred in the same stream. To ensure that the own name retained its unique familiar status, prior to participation in the study, participants were asked to list their full name as well as names of their closest family members and friends, including middle names and nicknames. These names were excluded entirely when creating the personalized streams for each participant.

### Experimental Procedure

In each trial, a Narrative Stream and a Barista Stream were presented concurrently. The Narrative Stream was presented with equal intensity to both ears and thus was perceived as originating from a central location. The Barista Stream was presented only to the right ear at a volume of 0.8 relative to the Narrative Stream. This created the perception of spatial separation between the streams, intended to simulate listening to a conversational partner sitting across from you, and hearing orders being called out from a peripheral location on the right (Figure 1B).

Participants were instructed to listen to <u>both</u> streams and perform a dual-task. For the Narrative Stream, they were instructed to follow the content of the narrative and answer four multiple-choice questions at the end of each trial. The questions could refer to specific details in the narrative (e.g. “Who did the speaker play with?”; answers: his brother, sister, cousin, friend), or to general themes in the narrative (e.g. “Why was the speaker sad?”). The second task was to monitor the Barista Stream, detect when a prescribed target name was mentioned in an order, and respond via button press as quickly as possible. The target name in each trial was either the participant’s own name or the control name, and clear instructions were given before each trial as to which target to detect. In addition, to ensure that participants knew exactly which name to listen for, a recording of the target name was played before the start of the trial, which participants could replay until they indicated they were ready to begin. At the end of the trial, participants received feedback regarding both tasks: their accuracy on each comprehension question and the percent of targets they had correctly detected (Figure 1A). The pairing of each Narrative Stream to an own name vs. control name trial was counterbalanced across participants.

After completing the main task, participants performed an additional task to assess their working memory capacity. We used a Hebrew adaptation of the Operation Span Working memory task (OSPAN), which is based on Shortened Version of the working memory task by Foster et al., (2014) (adaptation to Hebrew: https://englelab.gatech.edu/translatedtasks.html#hebrew). This task required the participants to solve simple arithmetic problems while also remembering a series of Hebrew letters. Each trial consisted of several (3-7) arithmetic problems each followed by a single letter of the Hebrew alphabet. At the end of each trial the participants were asked to recall the series of letters in the order that they were presented. Participants were instructed to maintain an average of at least 85% accuracy for the math equations throughout the task, and participants which achieved lower accuracy were excluded from the analysis. The test produces two main scores referring to the number of letters recalled and positioned correctly within the sequence (1) including only trials where all letters were correctly recalled (absolute OSPAN score) and (2) including all trials (partial OSPAN score). However, we used only the partial scores, as per recommendation of previous studies, since they make use of all available information (Friedman and Miyake, 2005; Redick et al., 2012; Đokić et al., 2018).

### Behavioral analysis

The behavioral accuracy on each task was analyzed separately in the own name and control name conditions. Accuracy on the comprehension questions about the Narrative Stream was the average number of correct answers in each trial (out of 4 questions). For the target detection task, a target was considered a hit if there was a button press within 1.5 seconds after the target. Otherwise, it was considered a miss. The percent of hits in each trial was calculated, as well as reaction times (RTs) for each target.

To test whether performance on either task was affected by the Target Type (own name vs. control) we performed a MANOVA analysis (Manova R function), with two dependent variables: Name Detection Accuracy and Narrative Comprehension Accuracy. The independent variables were Target Type (own name/control) and the number of targets per trial (1-3), and working-memory capacity (OSPAN-partial) was included as an individual factor.

We further tested whether there was a trade-off in performance on the two tasks, by calculating the Pearson correlation between accuracy on the narrative comprehension questions and target detection.

To complement these analyses of whole-trial-level responses, we also looked at the detection responses to individual targets and tested whether detection accuracy and RTs were different for own name vs. control name targets and added WM as an individual factor using a linear regression model (lmer R function).

### EEG Preprocessing

EEG preprocessing and analysis was performed using the matlab-based FieldTrip toolbox (Oostenveld et al., 2011) as well as custom written scripts. Raw data was first visually inspected and gross artifacts exceeding ±50 μV (that were not eye-movements) were removed. Independent Component Analysis (ICA) was performed to identify and remove components associated with horizontal or vertical eye movements as well as heartbeats. Any remaining noisy electrodes that exhibited either extreme high-frequency activity (>40Hz) or low-frequency activity/drifts (<1 Hz), were replaced with the weighted average of their neighbors using an interpolation procedure.

#### EEG Analysis - Speech Tracking

The first lens through which we analyzed the EEG data was to look at the speech tracking response to the two concurrent speech streams. One participant was removed from this analysis due to excessive artifacts, so this analysis was performed on 35 participants. This was done using the reverse correlation approach, as implemented in the mTRF MATLAB toolbox (Crosse et al., 2016). Briefly, in this analysis a Temporal Response Function (TRF) is estimated, which captures the (linear) relationship between the particular features of the stimulus (S) and the neural response (R) recorded while hearing it. In this case, the S represents the broadband envelope of the speech stimuli presented in each trial, which was extracted using an equally spaced filterbank between 100 to 10000 Hz based on Liberman’s cochlear frequency map. The narrowband filtered signals were summed across bands after taking the absolute value of the Hilbert transform for each one, resulting in a broadband envelope signal. The R was the continuous EEG data, after correcting for eye-movements (using ICA), and bandpass filtered between 0.8 and 20Hz (but without removing any additional artifacts). S and R were aligned in time, downsampled to 100Hz for computational efficiency and the first 1000ms of the data were removed to avoid effects of speech onset. TRF analysis was applied using both a decoding and encoding approach. For the decoding approach we used a multivariate approach, where the neural data from all EEG channels was used in order to reconstruct the envelope of the <u>two</u> auditory stimuli that were presented in each trial (S1 and S2; corresponding to the Narrative and Barista Streams). The advantage of the decoding approach is the simplicity of the results, which yields a single reconstruction accuracy for each stimulus, reflecting the Pearson’s correlation between the reconstructed S1* and S2* estimated by the model and the real S1 and S2.

However, the decoding model does not give insight in the spatio-temporal dynamics of the neural response itself. To this end, we used an encoding analysis which estimates a temporal response function (TRF) at each electrode, that captures the linear relationship between recorded EEG response (R) and each stimulus. Separate TRFs were estimated for the Narrative and Barista Speech, and the predictive power of each encoding model was assessed as the Pearson correlation between the predicted and actual neural response at each electrode.

Encoding and decoding models were run and optimized separately. Encoding TRFs were calculated over time lags ranging from −150 (pre-stimulus) to 450 msec, and the decoding analysis used time lags of −400 to 0 msec. Both models were optimized using a leave-one-out approach, where in each iteration, 39 trials are selected to train the model (train set), which was then used to predict either the neural response at each sensor (encoding) or the speech envelope of the two speakers (decoding) in the left-out trial (test set). The model’s goodness of fit (predictive power) is the Pearson correlation between the predicted and the actual signal. This procedure is repeated 40 times, with a different train-test partition in each iteration, and the predictive power is averaged over all 40 iterations. To prevent overfitting of the model, a ridge parameter (λ) was chosen as part of the cross-validation process. This parameter significantly influences the shape and amplitude of the TRF and therefore, rather than choosing a different λ for each participant (which would limit group-level analyses), a common λ value was selected for all participants. Specifically, for each participant, λ values ranging from 2^-5^ to 2^10^ were tested and the value yielding the average highest predictive power (across all channels\participants) was chosen as the common optimal λ. For both the encoding model and the decoding model this value was λ=2^9^. All results reported in here use these ridge parameters.

Statistical analysis of the decoding model was a pairwise t-test comparing the reconstruction accuracy of the Narrative Stream and the Barista Stream. For the encoding models, which were estimated separately for the Narrative and Barista Streams, statistical analysis focused on comparing the predictive power of each model to a null distribution, using a permutation test. This was done by randomly mismatching the S and R from different trials and estimating new encoding models for these mismatched pairs. This procedure was repeated 100 times for both the Narrative and Barista Speech, yielding a null distribution of predictive power values at each EEG channel and for each stimulus. We then z-scored the predictive power values from real-data set to this null-distribution, and the channels that exceeded an average of z>1.64 across participants (one-sided, p<0.05) were considered to have a significant speech tracking response to a particular stimulus.

#### EEG Analysis – RIDE ERP Analysis

For event-related analysis of the neural response to target names in the Barista Stream, the EEG data was segmented into epochs from 200ms before target onset to 1500ms after the target name onset. Only targets that were correctly detected (hits) were included in this analysis, and the number of remaining targets in each condition was randomly subsampled, to equate them across conditions. The segmented data was bandpass filtered between 0.5 to 13Hz and baseline corrected using the 200ms prior to target onset.

Since the target words were of prolonged and varied lengths, ranging from 500-750ms, and had different temporal profiles (van der Wulp, 2021), the classic method for deriving event-related potentials (ERPs) through simple averaging, is not appropriate. Instead, we applied Residue Iteration Decomposition (RIDE) analysis (http://cns.hkbu.edu.hk/RIDE.htm) (Ouyang et al., 2011), which accounts for latency differences across trials when extracting the average neural response. We chose this method because it recognizes that the recorded neural response is comprised of contributions from different subprocesses (e.g., stimulus-locked, reaction-locked), each of which may have different (and potentially jittered) latencies. The matlab-based RIDE toolbox (Ouyang et al., 2011) was used to decompose individual trials into stimulus-related (S), component-related (C), and response-related (R) components, and reconstruct a latency corrected RIDE-ERP. The time-windows defined for each component were as follows: The stimulus-related (S) component was defined between 0-400ms; two C-components – C1 and C2 - were defined, based on visual inspection of the raw ERP grand-average (prior to RIDE analysis, Figure 4A), with C1 spanning between 100-700ms, and C2-spanning from 400-1100ms. Last, a response-related R-component was defined, between −300 to 300ms relative to the reaction time measured for each target. This analysis was applied separately to responses to the own name and control name, and reconstructed RIDE-ERPs to each stimulus were extracted for each participant.

Group-level analysis of the RIDE-ERP responses also needs to take into account possible latency differences between participants (e.g. due to differences in the length of specific names). Hence, we subjected the RIDE-ERPs from each participant to a secondary RIDE analysis, using the same components and time-windows as in the individual analysis, as proposed by (Heidlmayr et al., 2021). This yielded a group-wise RIDE-ERP at each electrode, as well as the latencies by which each individual component (S, C1, C2 and R) was shifted when aggregating across participants. These latency-shifts were used to identify the prominent peaks in individual RIDE-ERPs and assess their amplitude, for the purpose of statistical testing. Our statistical analysis focuses only on the C1 and C2 components, that are clearly visible in the group-level RIDE response as a negative peak between 300-500ms and positive component between 500ms-700ms (Figure 4A).

Statistical analysis focused on comparing the amplitude of the C1 and C2 component in response to the own name vs. control name targets. A t-test was performed at each individual electrode on the amplitude of each component and corrected for multiple comparisons using a cluster-based permutation (TFCE correction, p<0.05; one-tailed).

#### GSR Analysis

Analysis of the GSR signal was performed using the matlab-based Ledalab toolbox (Benedek and Kaernbach, 2010), as well as custom written scripts. The raw data was manually inspected for distinguishable artifacts, which were fixed using linear interpolation. Then a continuous decomposition analysis (CDA) was performed on the full GSR signal and the time-locked response to individual target names was extracted in windows between 0 and 5 seconds from the onset of each target. GSR responses to target names were averaged across trials, separately for the own name and control name trials, but only targets that were correctly detected (hits) were included in the analysis. The number of remaining targets in each condition was randomly subsampled, to equate them across conditions. Statistical analysis focused on the average GSR amplitude between 2-3 seconds after the onset, which corresponds to the peak response. A one-way paired-wise t-test was performed comparing the GSR response to the own name vs the control name (JASP-Team, 2021), since previous findings suggest an a-priori larger GSR responses in the own name condition.

## Results

### Behavioral Results

#### Dual Task Performance

Performance on both tasks was quite good, with high accuracy on the Name Detection task (mean=92% hits, SD=5) and on the comprehension task (mean=80% correct, SD=7; Figure 2A). Furthermore, we observed no positive correlation nor tradeoffs between the two tasks (R= −0.09, p=0.62; Figure 2B). This is in line with participants’ self-evaluation post-experiment, in which most participants stated that they felt that they were able to perform both tasks pretty well. Note that task-difficulty and chance-level of the two tasks were not equated, therefore the accuracy values themselves are not quantitively comparable. Working memory capacity was only a marginally significant factor as found by the omnibus MANOVA analysis (F(1)=2.457=0.086) and was not found to be significantly correlated with any of our dependent variables (correlation with comprehension accuracy, r=0.14, p=0.41; target detection accuracy r=0.11, p=0.52; and target detection reaction time r=0.06, p=0.75; Figure 2C),

**Figure 2:**
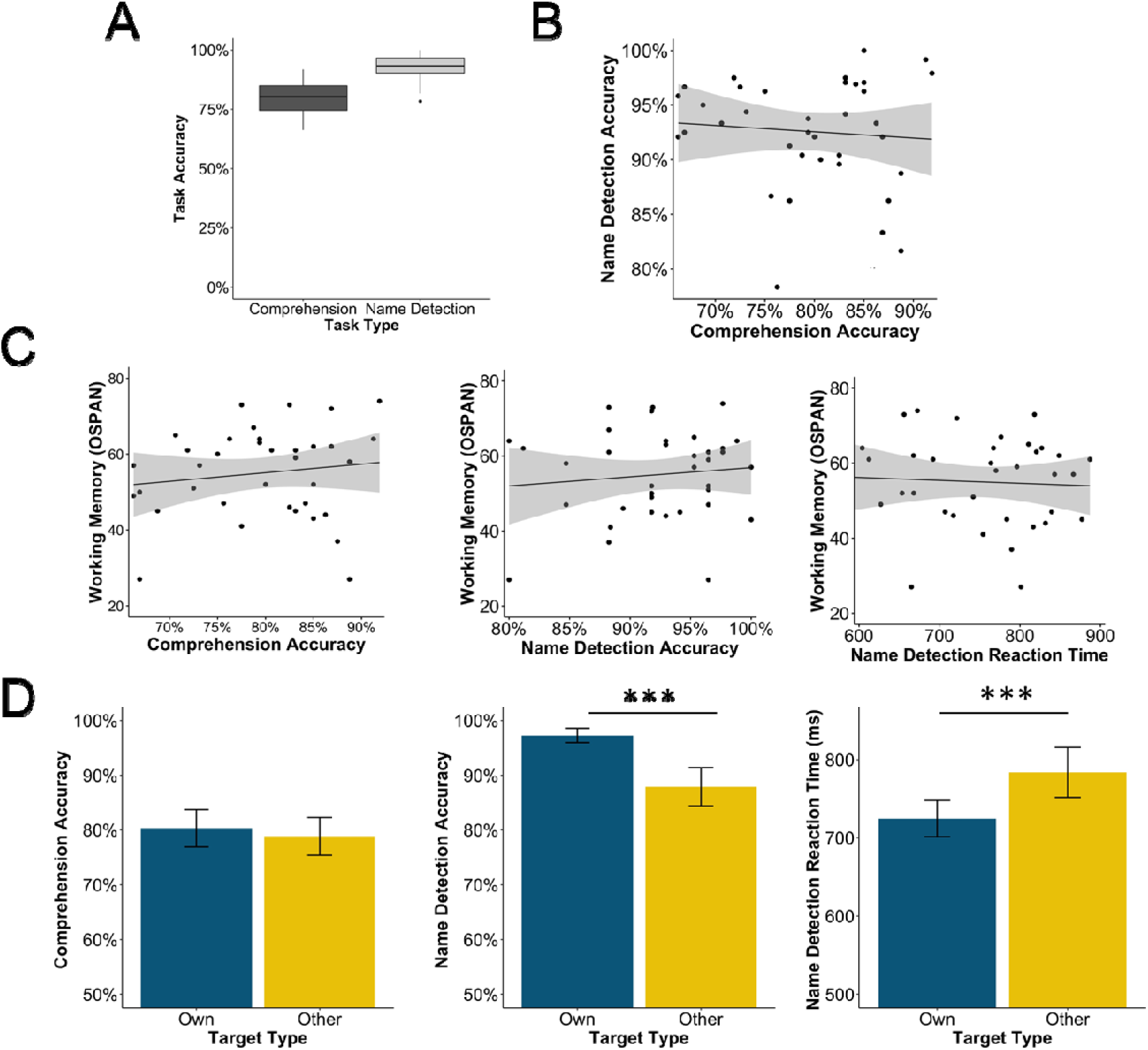
Behavioral Results: A) Box-plot showing the accuracy rates in the Narrative Comprehension and Name Detection tasks. B) Scatter plot of the dual-task accuracy shows no correlation or tradeoff between performance on the two tasks. The grey shaded area represents a 95% confidence interval. C) Scatter plots depicting the lack of a correlation between working memory capacity (OSPAN score) and our three behavioral outcomes: Narrative Comprehension Accuracy, Name Detection Accuracy and Name Detection Reaction Time. The grey shaded area represents a 95% confidence interval. D) Bar graphs showing group averages of Narrative Comprehension Accuracy, Name Detection Accuracy and Name Detection Reaction Time, comparing the own and control name condition. Error-bars indicate SEM. Asterisks *** represent significance at a level of p<=0.001.

#### Own Name Advantage

MANOVA testing of the effect of Target Type on accuracy, using both tasks (Narrative Comprehension and Name Detection) as dependent variables, revealed a significant main effect of Target Type (F(1)=44.08, p<0.001). When separating the analysis of each task into different ANOVAs we found that the main effects of Target Type and Target Number were only significant in the Name Detection Task, with higher detection rates of ones’ own name vs. the control, however no significant differences on the Narrative Comprehension Task was found between trials where participants were listening for their own name vs. the control name (Figure 2D). The linear regression analysis for the reaction times to individual targets also revealed a main effect of Target Type (F(1)=63.42, p=0.001), with faster reaction times to own name targets vs. control name targets (Figure 2D)

### EEG Results

#### Speech tracking analysis

We examined the neural representation of our two speech streams: the Narrative Stream and the Barista Stream, using both an encoding and decoding model. For the decoding model we used a multivariate model which included all EEG channels to reconstruct each of the speech stimuli envelopes in each trial and produced a value of reconstruction accuracy for each stimulus. Reconstruction accuracy was significantly higher for the Narrative Stream than for the Barista Stream [t(34)=10.99, p<0.001)], as shown in figure 3. In fact, the reconstruction accuracy for the Barista Stream was extremely low (r=0.0008).

**Figure 3:**
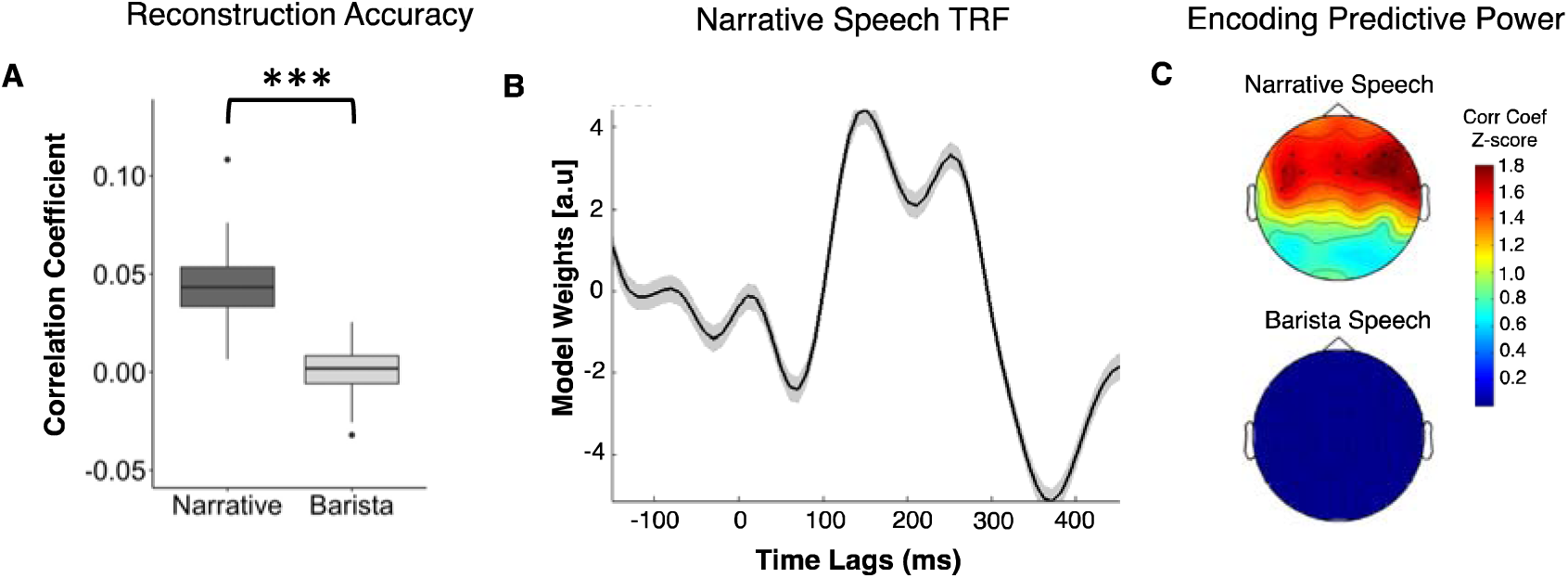
Speech tracking results: A) Reconstruction accuracy of the two speech stimuli across all participants using the decoding approach. The asterisks *** represent a significance difference between the stimuli (p<0.001). B) The TRF estimated for the Narrative Stream, averaged across participants, from the electrode with the highest normalized predictive power (FC6). Shaded error bars indicated SEM across participants. C) Scalp distribution the predictive power of the encoding mode, for both the Narrative and Barista Speech (z-scored relative to a null distribution). Asterisks indicate the electrodes where predictive power was significantly above chance (z>1.64, one tailed).

The encoding analysis was performed separately for the Narrative and Barista Streams (using a multivariate approach, where both stimuli are included in the same encoding model, did not change the results substantially). Corroborating the decoding results, here too we found a robust speech-tracking response to the Narrative Stream, which had significant predictive power in a cluster of 15 central electrodes (Figure 3B&C). However, the predictive power of the Barista Stream TRF was extremely low and did not reach significance, relative to the null distribution (Figure 3C).

### Response to Target Names in Barista Stream

Even though no reliable neural speech tracking response was obtained for the Barista Stream, we do observe robust time-locked responses to the target-words in this stream. Figure 4B shows the group-level RIDE ERP, as well as the classic ERP grand-average (before RIDE analysis; Figure 4A). In both time-courses, two main peaks are visible in response to both the own name and the control name targets – a negative peak between 300-500ms, and a positive peak between 500-700ms. These were used to define the C1 and C2 components of the RIDE analysis, which was used to correct for jitter across participants and trials. Statistical comparison of the peak-amplitude of the resulting RIDE components showed that both were significantly larger in response to the own name target vs. the control name. The negative peak, maximal at 400ms, had a centro-frontal scalp distribution and was significantly more negative in all 64 electrodes (p<0.001, TFCE cluster corrected). The later positive peak, maximal at 600ms, showed a more posterior scalp-distribution and was significantly modulated by Target Type in a cluster of 11 posterior electrodes (p<0.05, TFCE cluster corrected).

**Figure 4:**
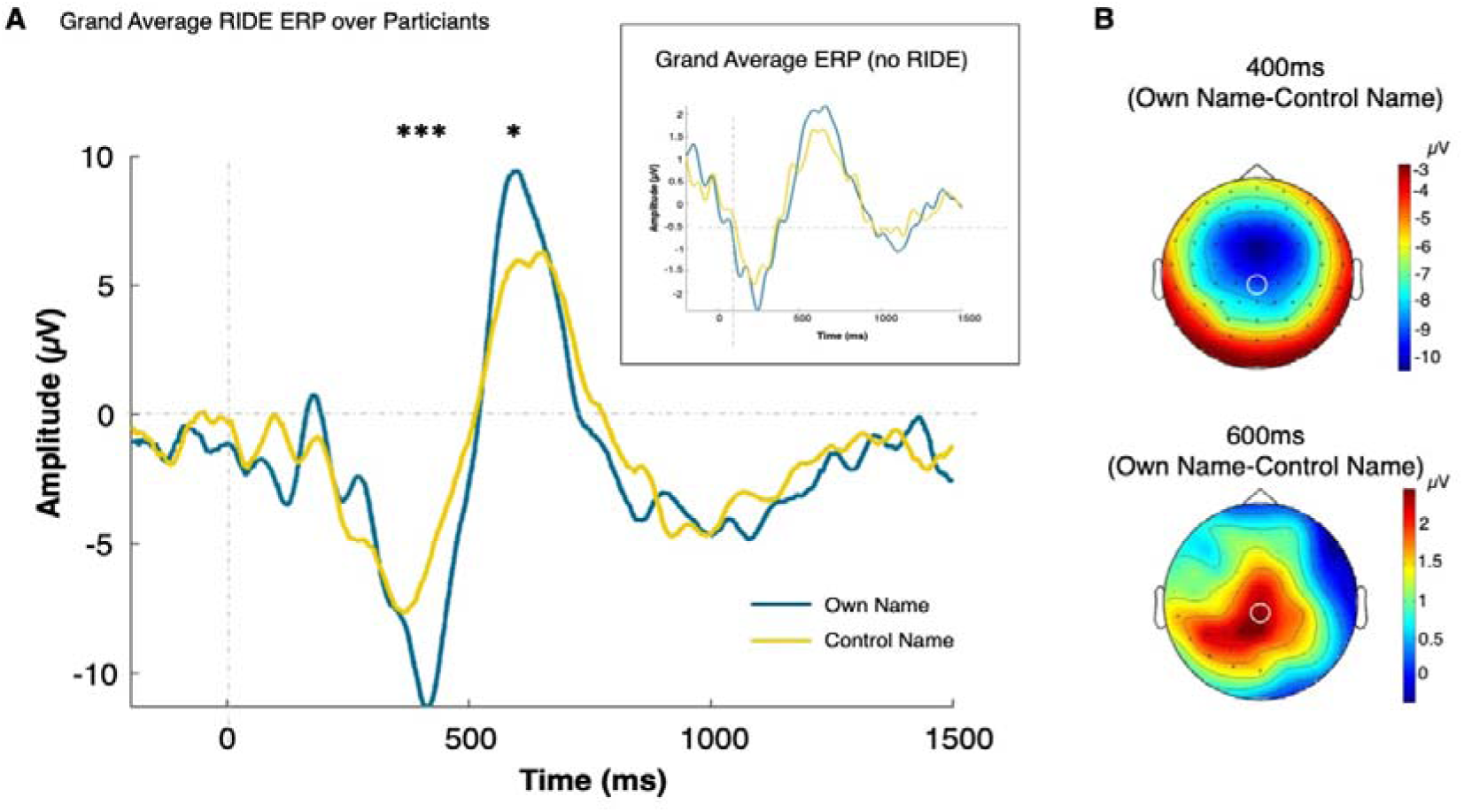
A. Group-level grand average RIDE-ERP in response to hearing the own name and control name targets, shown from the central electrode Pz (indicated on the topographies in panel B). The asterisks represent significant differences (*=0.05, **=0.01, ***=0.001). Inset: The classic grand average ERP for these stimuli, before application of the RIDE analysis, which was used to define the time-windows for estimation of the two RIDE components. B. Scalp-topographies showing the differences between responses to hearing the own name vs. control name targets, at the mid-latency negative peak (400ms) and the later positive peak (600ms). Electrodes where significant differences were found are indicated with black asterisks. Electrode Pz, for which the time courses in A are shown, is marked with a white circle.

### GSR Results

The event-related GSR response was extracted in response to hearing ones’ own name and the control target. The response peaked between 2-3 seconds after stimulus onset and was significantly larger after hearing ones’ own name vs. the control name (t(35)=3.152, p=0.002; Figure 5).

**Figure 5:**
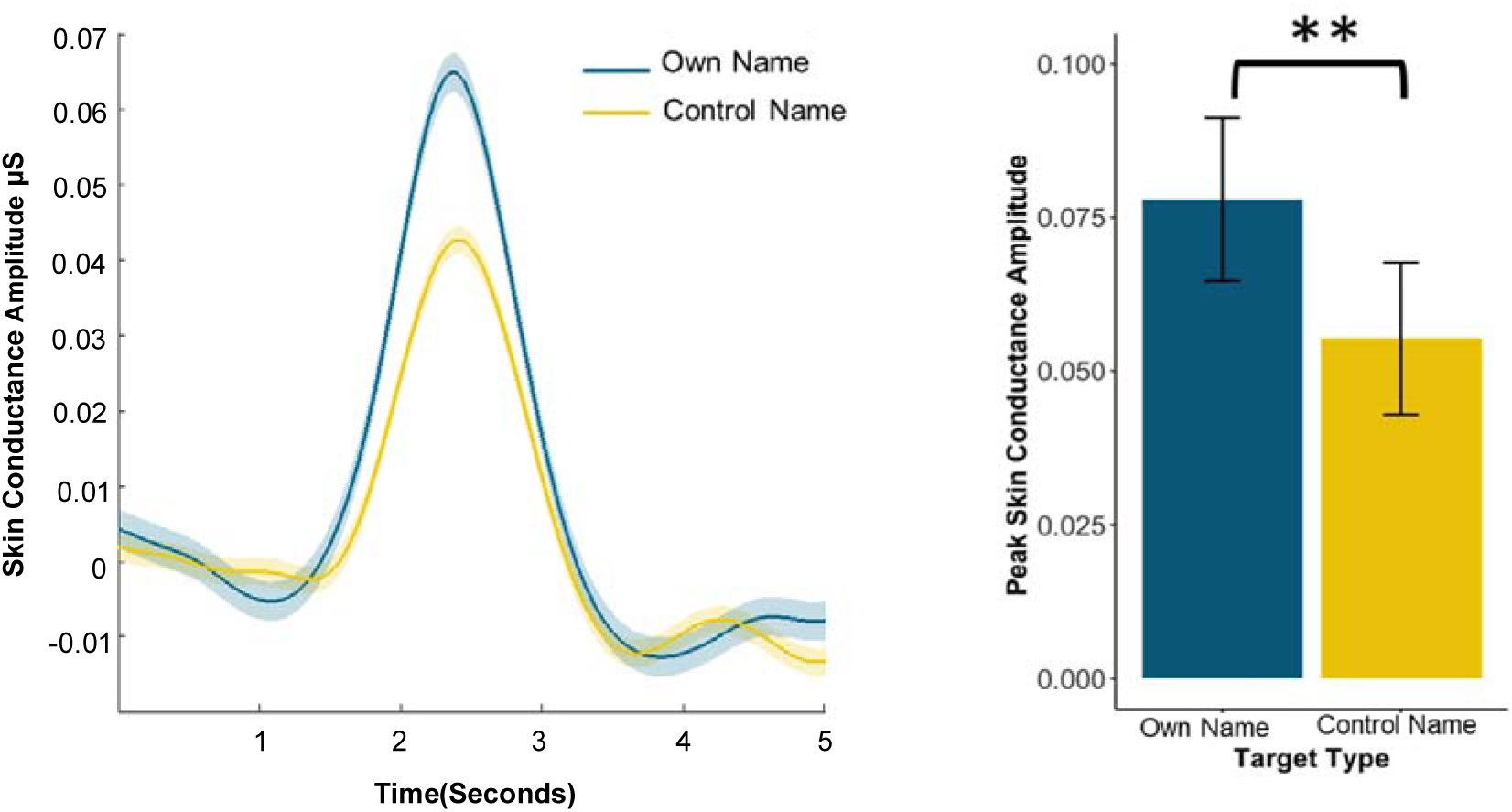
Skin Conductance (GSR) results comparing the own and control name condition. Left: Groups-level event related skin conductance response to both name targets. Right: Bar graph representing the average skin conductance levels around the peak, for both target name conditions, which was significantly larger in response to hearing ones’ own name. Error bars indicate SEM. Asterisks ** represent significance at a level of p<0.01.

## Discussion

The dual-task paradigm used here mimics real-life situations in which a listener tries to pay attention to one speaker and understand what they are saying, and at the same time also monitors another speech stream for potentially relevant information (in this case, a target name). This task-combination was chosen for its ecological relevance, since many real-life situations involve distributed, yet imbalanced, attention towards concurrent stimuli. In that vein, we chose to use highly contextual speech materials for both streams, as opposed to the arbitrary word-lists or brief highly-structured sentences used in previous studies of attention to concurrent speech (Berlad and Pratt, 1995; Wood and Cowan, 1995; Brungart et al., 2001, 2005; Conway et al., 2001; Perrin et al., 2005; Best et al., 2005, 2006, 2010; Shafiro and Gygi, 2007; Colflesh and Conway, 2007; Abel et al., 2012; Tateuchi et al., 2012; Naveh-Benjamin et al., 2014; Gygi and Shafiro, 2014). Moreover, by imposing explicit behavioral demands with regard to <u>both</u> streams, the dual-task design can help further our understanding of how the system toggles the need to process information from competing sources, circumventing the ‘ground-truth’ ambiguity of selective attention studies. In discussing the current results, it is helpful to bear in mind that they pertain only to the specific task-combination used here. In order to gain a deeper understanding of the capacity for processing simultaneous speech, additional research is required using task-combinations that impose varied depth of processing requirements on the listener (Mattys et al., 2009; Vazquez Alvarez and Brewster, 2010; Xia et al., 2015; Murphy et al., 2017; Fazal et al., 2022).

### Processing concurrent speech – Neural prioritization without behavioral tradeoffs

Analysis of the behavioral and neural responses to the two concurrent speech streams reveals an interesting pattern. Participants achieved highly accurate performance on both the narrative comprehension questions and target detection task in the Barista Stream, with no apparent behavioral trade-off between them, indicating that this combination of tasks did not overload participants’ cognitive capacity. Moreover, adequate performance on this dual-task was not correlated with individual working-memory capacity (WMC), further supporting its relative low-cognitive-demand nature (Conway et al., 2001; Colflesh and Conway, 2007; Gygi and Shafiro, 2012; Naveh-Benjamin et al., 2014). At the same time, the neural speech tracking analysis shows that the Narrative Stream was represented more robustly than the Barista Stream, a pattern reminiscent of the enhanced speech-tracking of task-relevant speech in selective attention studies (Kerlin et al., 2010; Ding and Simon, 2012b, 2012a; Mesgarani and Chang, 2012; Power et al., 2012; Zion Golumbic et al., 2013; O’Sullivan et al., 2015; Fuglsang et al., 2017; Fiedler et al., 2019; Har-shai Yahav and Zion Golumbic, 2021; Kaufman and Golumbic, 2022). In discussing this result, we acknowledge the possibility that the lack of a reliable speech-tracking response to the Barista Stream may be due, at least in part, to the highly structured nature of this stimulus which contains substantial autocorrelation. The inverse-correlation approach used to estimate the TRFs is highly affected by the degree of autocorrelation in the stimulus, and hence the two speech stimuli are not directly comparable (Crosse et al., 2016). Nonetheless, even highly rhythmic stimuli have been shown to produce reliable TRFs in other studies (Leahy et al., 2021; Lerousseau et al., 2021), therefore, this methodological caveat alone probably does not account for the lack of any reliable signature of neural tracking of the Barista Stream observed here.

The apparent discrepancy between the behavioral and neural results reminds us that different listening strategies can be employed to achieve good behavioral outcomes. Given the different task-demands for the two streams, listeners may have employed different ways of listening to each one, so as to optimize performance without overly taxing the system. One possibility, that is consistent with the observed speech-tracking pattern, is that the Narrative Stream was tracked continuously since its task required understanding its entire content, whereas detecting target words in the Barista Stream was done by applying an acoustic template-matching strategy, but the Barista Stream was not represented in full. Supporting this possibility, previous studies have shown individual words can be detected under noisy conditions using a low-level and possibly automatic (Shiffrin and Schneider, 1977; Logan, 1988) mechanism of temporal ‘glimpsing’, that requires only a relatively sparse spectro-temporal representation of the speech (Assmann and Summerfield, 1989; Cooke, 2006; Heald and Nusbaum, 2014; Best et al., 2016; Josupeit et al., 2016; Josupeit and Hohmann, 2017; Kidd, 2017; Daube et al., 2019). Hence, it is possible that the good performance achieved on the Target Detection Task simply did not require full speech tracking of the Barista Stream, leading to observed preferential neural tracking of the Narrative Stream.

A similar dissociation between behavioral performance and neural tracking of concurrent speech was observed in a previous study of distributed attention (Kaufman & Zion Golumbic). In that study participants were required to listen to the content of two Narrative Streams and answer comprehension questions about <u>both</u> of them. There, we observed a tradeoff in the neural speech tracking of the two speakers which were not simultaneously represented to the same degree, even though they were equally task-relevance, potentially indicating inherent limitation for fully tracking two speakers at once.

And yet, despite their imbalanced neural representation, behavioral performance was similarly good for both speakers, and reduced neural tracking was not associated with lower comprehension levels. Together with the current findings, these results serve as an important reminder of the non-binary nature of attention to speech, namely that speech is not merely “attended” or “unattended”, but rather that there are different ways of attending to speech and neural responses to speech should be interpreted with regard to the specific task participants were given and how well they performed it.

The current results also emphasize an important limitation of the speech-tracking analysis approach. This approach has been extremely influential in advancing speech-processing research into the realm of continuous, naturalistic speech (Simon, 2015; Brodbeck and Simon, 2020). At the same time, it clearly does not capture all aspects of the neural response to speech processing. Specifically, observing a diminished, or unreliable, speech-tracking response to a particular speaker (e.g., for the Barista Stream in the current study) does not necessarily imply that the speech was not processed by the listener, as is evident here from the robust behavioral, physiological, and neural response to target words in the Barista Stream.

### Own Name Advantage

This unique design of this study allowed us to revisit the notion of an advantage for detecting ones’ own name, under ecological circumstances that mimic the type of real-life situations where this advantage might manifest. First reported by Cherry (1953), the own name advantage has been demonstrated in a variety of behavioral and neural studies, arguably reflecting its ‘special status’ in the perceptual system (Deutsch and Deutsch, 1963; Gallagher, 2000; Perrin et al., 2005; Alho and Vorobyev, 2007). The most commonly known, yet also somewhat controversial, finding is of detecting ones’ name in a presumed “unattended” stream, in selective attention paradigm (Wood and Cowan, 1995; Röer et al., 2013; Ljungberg et al., 2014; Naveh-Benjamin et al., 2014; Holtze et al., 2021) although behavioral advantages have been shown in a variety of other paradigms (Perrin et al., 1999; Tamura et al., 2012; Tateuchi et al., 2012; Röer et al., 2013; Lechinger et al., 2016; Liu et al., 2019; Jijomon and Vinod, 2021; Ye et al., 2021). Several ERP components have also been shown to be enhanced in response to hearing or reading ones’ own name vs. other names or words. These include the central P300 response, that is associated with target detection (Berlad and Pratt, 1995; Folmer and Yingling, 1997; Perrin et al., 1999, 2005; Tacikowski and Nowicka, 2010; Tamura et al., 2012; Holtze et al., 2021; Lu et al., 2021) the central N400 response associated with semantic processing (Müller and Kutas, 1997), and the parietal P600 response (Berlad and Pratt, 1995; Eichenlaub et al., 2012), which is also linked to semantic processing and memory retrieval. However, almost all previous studies of the own name advantage have done so using highly artificial stimuli where names were presented as part of word-lists rather than embedded in ecologically-relevant contexts.

The current results confirm the ecological validity of this body of literature, by showing a robust advantage to hearing and responding to ones’ own name in a situation simulating how this might manifest in real-life. We show faster and more accurate responses to ones’ name, which was accompanied by higher GSR responses representing a heightened state of arousal and alertness (Gronau et al., 2003). The novel RIDE-ERP analysis employed here also revealed an enhanced neural response to hearing ones’ name, which is qualitatively similar to previously reported ERP effects. Drawing direct associations between known ERP components and the two RIDE-ERP peaks observed here (the mid-latency negativity and a late positivity) is beyond the scope of the current study, particularly since the RIDE analysis essentially shifts the latency of these peaks around (Ouyang et al., 2011, 2013, 2015a, 2015b, 2016).

At the same time, the temporal morphology and topographical distributions of these responses could be analogous to at least two known ERP effects of responses ones’ name: (1) the N2-P3 complex, ERP components associated with target detection that have been shown to be enhanced in response for ones’ own name in more controlled experiments (Folmer and Yingling, 1997; Perrin et al., 2005; Tamura et al., 2012; Liu et al., 2019), or (2) the later N400-P600 responses that are linked to processing and memory retrieval and have also been reported as sensitive to hearing ones’ name (Berlad and Pratt, 1995; Müller and Kutas, 1997; Eichenlaub et al., 2012; Tamura et al., 2012). Additional research and replication is necessary to link the RIDE-ERP effects observed here to the more well-studied ERP responses and deepen our mechanistic understanding on the neural processes generating the observed heightened response to ones’ own name. Nonetheless, the combined behavioral, physiological and neural responses reported here provide strong support for the ‘special status’ of one’s own name, which may draw more attention in real-life multi-speaker contexts, relative to non-personal words.

### Implication for Selective Attention studies

One of the motivations for designing this dual-task experiment, was the hope that it might inform ongoing conversations about whether or not words in so-called “unattended” speech streams can be processed linguistically (Rivenez et al., 2006, 2008; Carey et al., 2014; Aydelott et al., 2015b; Tóth et al., 2019; Har-shai Yahav and Zion Golumbic, 2021). As the current results clearly show, when listeners actively try to glean information from two concurrent speech streams (for comprehension or word-detection purposes), this is quite possible and does not excessively tax their cognitive resources. Admittedly, here participants knew in advance which target-words to expect, which likely made this task easier for them. However, these results are in line with other studies showing that people can distribute their attention among two speech stimuli and accurately report content from both, even without prior expectations (Shinn-Cunningham and Ihlefeld, 2004; Shafiro and Gygi, 2007; Ihlefeld and Shinn-Cunningham, 2008; Gygi and Shafiro, 2014; Lambez et al., 2020; Kaufman and Golumbic, 2022). Given that distributing attention does not seem to bear considerable costs to performance, it seems likely that listeners might employ a distributed listening strategy in selective attention paradigms as well and monitor task-irrelevant speech even when not instructed to. This explanation would easily explain a wealth of findings regrading processing irrelevant speech, including the detection of ones’ own name (Conway et al., 2001; Dupoux et al., 2003; Rivenez et al., 2006, 2007, 2008; Best et al., 2010; Iyer et al., 2013; Röer et al., 2013; Naveh-Benjamin et al., 2014; Aydelott et al., 2015; Kidd and Arciuli, 2016; Röer et al., 2017; Har-shai Yahav and Zion Golumbic, 2021; Holtze et al., 2021; Kaufman and Golumbic, 2022). Considering the ‘ground-truth’ uncertainty regarding what listeners’ actually do in selective attention paradigms, and the relatively simple tasks they are asked to perform, this is not a possibility we can easily rule out. It is, of course, possible that increasing the task difficulty or level of engagement with one speech stream would reduce the available resources for processing a second one, in line with ideas from load-theory of attention (Mattys et al., 2009; Murphy et al., 2017). However, the impact of perceptual and cognitive load on speech processing in multi-talker contexts and its interaction with the ability to process the content of a secondary speech stream has not, of yet, been studied in a sufficiently principled manner (Gagné et al., 2017; see comparison of results by Har-shai Yahav and Zion Golumbic, 2021 vs. Ding et al., 2016, for additional speculation regarding the effect of task difficulty on linguistic analysis of task-irrelevant speech).

## Conclusions

The current results broaden the ongoing conversation regarding the way our brains deal with the multitude of stimuli in our real-life environments. Rather than assuming that only one stimulus is of interest to listeners, this study recognizes that most real-life situations require some degree of multiplexing and processing of multiple stimuli. Importantly, our results demonstrate that individuals are highly capable of carrying out this multiplexing, at least under moderate conditions. As the field of attention research ventures into studying human performance and neural operations under increasingly ecological conditions (Parsons, 2015; Shamay-Tsoory and Mendelsohn, 2019; Hölle et al., 2020; Holtze et al., 2021), it will become increasingly important to understand how attention is deployed dynamically to monitor concurrent stimuli and the capabilities and limitations of multiplexed listening.

## Acknowledgments

This work was supported by Israel Science Foundation grant # 2339/20.

## References

Abel SM, Nakashima A, Smith I (2012) Divided listening in noise in a mock-up of a military command post. Mil Med 177:436–443.

Agmon G, Yahav PH-S, Ben-Shachar M, Golumbic EZ (2021) Attention to speech: mapping distributed and selective attention systems. Cereb Cortex.

Alho K, Vorobyev VA (2007) Brain activity during selective listening to natural speech. Front Biosci 12:3167–3176.

Assmann PF, Summerfield Q (1989) Modeling the perception of concurrent vowels: vowels with the same fundamental frequency. J Acoust Soc Am 85:327–338.

Aydelott J, Jamaluddin Z, Nixon Pearce S (2015) Semantic processing of unattended speech in dichotic listening. J Acoust Soc Am 138:964–975.

Baldock J, Kapadia S, van Steenbrugge W (2019) The task-evoked pupil response in divided auditory attention tasks. J Am Acad Audiol 30:264–272.

Beaman CP (2004) The irrelevant sound phenomenon revisited: what role for working memory capacity? J Exp Psychol Learn Mem Cogn 30:1106–1118.

Benedek M, Kaernbach C (2010) A continuous measure of phasic electrodermal activity. J Neurosci Methods 190:80–91.

Berlad I, Pratt H (1995) P300 in response to the subject’s own name. Electroencephalogr Clin Neurophysiol 96:472–474.

Best V, Gallun FJ, Ihlefeld A, Shinn-Cunningham BG (2006) The influence of spatial separation on divided listening. J Acoust Soc Am 120:1506–1516.

Best V, Gallun FJ, Mason CR, Kidd G, Shinn-Cunningham BG (2010) The impact of noise and hearing loss on the processing of simultaneous sentences. Ear Hear 31:213–220.

Best V, Ihlefeld A, Shinn-Cunningham B (2005) The effect of auditory spatial layout in a divided attention task. Proc ICAD 5:17–22.

Best V, Keidser G, Buchholz JM, Freeston K (2016) Development and preliminary evaluation of a new test of ongoing speech comprehension. Int J Audiol 55:45–52.

Boersma P, Weenink D (2010) PRAAT software.

Boudewyn MA, Carter CS (2018) I must have missed that: alpha-band oscillations track attention to spoken language. Neuropsychologia 117:148–155.

Broadbent DE (1958) Perception and Communication. London: Pergamon Press.

Brodbeck C, Jiao A, Hong LE, Simon JZ (2020) Neural speech restoration at the cocktail party: auditory cortex recovers masked speech of both attended and ignored speakers Malmierca MS, ed. PLOS Biol 18:e3000883.

Brodbeck C, Simon JZ (2020) Continuous speech processing. Curr Opin Physiol 18:25–31.

Brungart DS, Iyer N (2012) Better-ear glimpsing efficiency with symmetrically-placed interfering talkers. J Acoust Soc Am 132:2545–2556.

Brungart DS, Kordik AJ, Simpson BD (2005) Audio and visual cues in a two-talker divided attention speech-monitoring task. Hum Factors J Hum Factors Ergon Soc 47:562–573.

Brungart DS, Simpson BD, Ericson MA, Scott KR (2001) Informational and energetic masking effects in the perception of multiple simultaneous talkers. J Acoust Soc Am 110:2527– 2538.

Carey D, Mercure E, Pizzioli F, Aydelott J (2014) Auditory semantic processing in dichotic listening: effects of competing speech, ear of presentation, and sentential bias on n400s to spoken words in context. Neuropsychologia 65:102–112.

Cherry CE (1953) Some experiments on the recognition of speech, with one and two ears. J Acoust Soc Am 25:975–979.

Colflesh GJH, Conway ARA (2007) Individual differences in working memory capacity and divided attention in dichotic listening. Psychon Bull Rev 14:699–703.

Conway ARA, Cowan N, Bunting MF (2001) The cocktail party phenomenon revisited: the importance of working memory capacity. Psychon Bull Rev 8:331–335.

Cooke M (2006) A glimpsing model of speech perception in noise. J Acoust Soc Am 119:1562–1573.

Crosse MJ, Di Liberto GM, Bednar A, Lalor EC (2016) The multivariate temporal response function (mtrf) toolbox: a matlab toolbox for relating neural signals to continuous stimuli. Front Hum Neurosci 10:604.

Dai B, McQueen JM, Terporten R, Hagoort P, Kosem A (2021) Distracting linguistic information impairs neural tracking of attended speech. bioRxiv 21.

Daube C, Ince RAA, Gross J (2019) Simple acoustic features can explain phoneme-based predictions of cortical responses to speech. Curr Biol 29:1924–1937.e9.

Del Giudice R, Lechinger J, Wislowska M, Heib DPJ, Hoedlmoser K, Schabus M (2014) Oscillatory brain responses to own names uttered by unfamiliar and familiar voices. Brain Res 1591:63–73.

Deutsch JA, Deutsch D (1963) Attention: some theoretical considerations. Psychol Rev 70:80–90.

Ding N, Melloni L, Zhang H, Tian X, Poeppel D (2016) Cortical tracking of hierarchical linguistic structures in connected speech. Nat Neurosci 19:158–164.

Ding N, Pan X, Luo C, Su N, Zhang W, Zhang J (2018) Attention is required for knowledge-based sequential grouping: insights from the integration of syllables into words. J Neurosci 38:1178–1188.

Ding N, Simon JZ (2012a) Neural coding of continuous speech in auditory cortex during monaural and dichotic listening. J Neurophysiol 107:78–89.

Ding N, Simon JZ (2012b) Emergence of neural encoding of auditory objects while listening to competing speakers. Proc Natl Acad Sci 109:11854–11859.

Đokić R, Koso-drljevi M, Apo N (2018) Working memory span tasks: group administration and omitting accuracy critereon do not change metric characteristics. PLoS One 13.

Driver J (2001) A selective review of selective attention research from the past century. Br J Psychol 92 Part 1:53–78.

Dupoux E, Kouider S, Mehler J (2003) Lexical access without attention? explorations using dichotic priming. J Exp Psychol Hum Percept Perform 29:172–184.

Eichenlaub JB, Ruby P, Morlet D (2012) What is the specificity of the response to the own first-name when presented as a novel in a passive oddball paradigm? an erp study. Brain Res 1447:65–78.

Fazal MA ul, Ferguson S, Saeed Z (2022) Investigating cognitive workload in concurrent speech-based information communication. Int J Hum Comput Stud 157:102728.

Fiedler L, Wöstmann M, Herbst SK, Obleser J (2019) Late cortical tracking of ignored speech facilitates neural selectivity in acoustically challenging conditions. Neuroimage 186:33– 42.

Folmer R, Yingling CD (1997) Auditory p3 responses to name stimuli. Brain Lang 311:306– 311.

Foster JL, Shipstead Z, Harrison TL, Hicks KL, Redick TS, Engle RW (2014) Shortened complex span tasks can reliably measure working memory capacity. Mem Cogn 43:226–236.

Friedman N, Miyake A (2005) Comparison of four scoring methods for the reading span test. Behav Res Methods 37:581–590.

Fuglsang SA, Dau T, Hjortkjær J (2017) Noise-robust cortical tracking of attended speech in real-world acoustic scenes. Neuroimage 156:435–444.

Gagné JP, Besser J, Lemke U (2017) Behavioral assessment of listening effort using a dual-task paradigm: a review. Trends Hear 21:1–25.

Gallagher S (2000) Philosophical conceptions of the self: implications for cognitive science. Trends Cogn Sci 4:14–21.

Gronau N, Cohen A, Ben-Shakhar G (2003) Dissociations of personally significant and task-relevant distractors inside and outside the focus of attention: a combined behavioral and psychophysiological study. J Exp Psychol Gen 132:512–529.

Gygi B, Shafiro V (2012) Spatial and temporal factors in a multitalker dual listening task. Acta Acust united with Acust 98:142–157.

Gygi B, Shafiro V (2014) Spatial and temporal modifications of multitalker speech can improve speech perception in older adults. Hear Res 310:76–86.

Har-shai Yahav P, Zion Golumbic E (2021) Linguistic processing of task-irrelevant speech at a cocktail party. Elife 10:e6509.

Heald SLM, Nusbaum HC (2014) Speech perception as an active cognitive process. Front Syst Neurosci 8:1–15.

Heidlmayr K, Ferragne E, Isel F (2021) Neuroplasticity in the phonological system: the pmn and the n400 as markers for the perception of non-native phonemic contrasts by late second language learners. Neuropsychologia 156.

Holender D (1986) Semantic activation without conscious identification in dichotic listening, parafoveal vision, and visual masking: a survey and appraisal. Behav Brain Sci 9:1–23.

Hölle D, Meekes J, Bleichner MG (2020) Mobile ear-eeg to study auditory attention in everyday life. bioRxiv:2020.09.09.287490.

Höller Y, Kronbichler M, Bergmann J, Crone JS, Ladurner G, Golaszewski S (2011) EEG frequency analysis of responses to the own-name stimulus. Clin Neurophysiol 122:99– 106.

Holtze B, Jaeger M, Debener S, Adiloğlu K, Mirkovic B (2021) Are they calling my name? attention capture is reflected in the neural tracking of attended and ignored speech. Front Neurosci 15.

Humes LE, Lee JH, Coughlin MP (2006) Auditory measures of selective and divided attention in young and older adults using single-talker competition. J Acoust Soc Am 120:2926– 2937.

Ihlefeld A, Shinn-Cunningham B (2008) Disentangling the effects of spatial cues on selection and formation of auditory objects. J Acoust Soc Am 124:2224–2235.

Iyer N, Thompson ER, Simpson BD, Brungart D, Summers V (2013) Exploring auditory gist: comprehension of two dichotic, simultaneously presented stories. Proc Meet Acoust 19.

JASP-Team (2021) JASP (version 0.15).

Jijomon CM, Vinod AP (2021) Detection and classification of long-latency own-name auditory evoked potential from electroencephalogram. Biomed Signal Process Control 68:102724.

Josupeit A, Hohmann V (2017) Modeling speech localization, talker identification, and word recognition in a multi-talker setting. J Acoust Soc Am 142:35–54.

Josupeit A, Kopco N, Hohmann V (2016) Modeling of speech localization in a multi-talker mixture using periodicity and energy-based auditory features. J Acoust Soc Am 139:2911–2923.

Kahneman D, Treisman AM (1984) Changing views of attention and automaticity. Var Atten:29–61.

Kaufman M, Golumbic EZ (2022) Capacity and tradeoffs in neural encoding of concurrent speech during selective and distributed attention. bioRxiv:2022.02.08.479628.

Kerlin JR, Shahin AJ, Miller LM (2010) Attentional gain control of ongoing cortical speech representations in a “cocktail party.” J Neurosci 30:620–628.

Kidd E, Arciuli J (2016) Individual differences in statistical learning predict children’s comprehension of syntax. Child Dev 87:184–193.

Kidd G (2017) Enhancing auditory selective attention using a visually guided hearing aid. J Speech Lang Hear Res 60:3027.

Koelewijn T, Shinn-Cunningham BG, Zekveld AA, Kramer SE (2014) The pupil response is sensitive to divided attention during speech processing. Hear Res 312:114–120.

Lachter J, Forster KI, Ruthruff E (2004) Forty-five years after broadbent (1958): still no identification without attention. 111:880–913.

Lambez B, Agmon G, Har-Shai Yahav P, Rassovsky Y, Zion Golumbic E (2020) Paying attention to speech: the role of working memory capacity and professional experience. Attention, Perception, Psychophys 82:3594–3605.

Lane DM, Pearson DA (1982) The development of selective attention. Merrill Palmer Q 28:317–337.

Leahy J, Kim SG, Wan J, Overath T (2021) An analytical framework of tonal and rhythmic hierarchy in natural music using the multivariate temporal response function. Front Neurosci 15:1–13.

Lechinger J, Wielek T, Blume C, Pichler G, Michitsch G, Donis J, Gruber W, Schabus M (2016) Event-related eeg power modulations and phase connectivity indicate the focus of attention in an auditory own name paradigm. J Neurol 263:1530–1543.

Lerousseau JP, Trébuchon A, Morillon B, Schön D (2021) Frequency selectivity of persistent cortical oscillatory responses to auditory rhythmic stimulation. J Neurosci 41:7991– 8006.

Liu L, Li W, Li J, Lou L, Chen J (2019) Temporal features of psychological and physical self-representation: an erp study. Front Psychol 10.

Ljungberg JK, Parmentier FBR, Jones DM, Marsja E, Neely G (2014) “What’s in a name?” “no more than when it’s mine own”. evidence from auditory oddball distraction. Acta Psychol (Amst) 150:161–166.

Logan GD (1988) Toward an instance theory of automatization. Psychol Rev 95:492–527.

Lu L, Sheng J, Liu Z, Gao JH (2021) Neural representations of imagined speech revealed by frequency-tagged magnetoencephalography responses. Neuroimage 229:117724.

Mattys SL, Brooks J, Cooke M (2009) Recognizing speech under a processing load: dissociating energetic from informational factors. Cogn Psychol 59:203–243.

Mesgarani N, Chang EF (2012) Selective cortical representation of attended speaker in multi-talker speech perception. Nature 485:233–236.

Moray N (1959) Attention in dichotic listening: affective cues and the influence of instructions. Q J Exp Psychol 11:56–60.

Müller HM, Kutas M (1997) What’s in a name? electrophysiological differences between spoken nouns, proper names and one’s won name. Neuroreport 8:221–225.

Murphy S, Spence C, Dalton P (2017) Auditory perceptual load: a review. Hear Res 352:40– 48.

Näätänen R (1988) Implication of erp data for psychological theories of attention. Biol Psychol 26:117–163.

Naveh-Benjamin M, Kilb A, Maddox GB, Thomas J, Fine HC, Chen T, Cowan N (2014) Older adults do not notice their names: a new twist to a classic attention task. J Exp Psychol Learn Mem Cogn 40:1540–1550.

O’Sullivan JA, Power AJ, Mesgarani N, Rajaram S, Foxe JJ, Shinn-Cunningham BG, Slaney M, Shamma SA, Lalor EC (2015) Attentional selection in a cocktail party environment can be decoded from single-trial eeg. Cereb Cortex 25:1697–1706.

Oostenveld R, Fries P, Maris E, Schoffelen J-M (2011) FieldTrip: open source software for advanced analysis of meg, eeg, and invasive electrophysiological data. Comput Intell Neurosci 2011.

Ouyang G, Herzmann G, Zhou C, Sommer W (2011) Residue iteration decomposition (ride): a new method to separate erp components on the basis of latency variability in single trials. Psychophysiology 48:1631–1647.

Ouyang G, Schacht A, Zhou C, Sommer W (2013) Overcoming limitations of the erp method with residue iteration decomposition (ride): a demonstration in go/no-go experiments. Psychophysiology 50:253–265.

Ouyang G, Sommer W, Zhou C (2015a) A toolbox for residue iteration decomposition (ride)-a method for the decomposition, reconstruction, and single trial analysis of event related potentials. J Neurosci Methods 250:7–21.

Ouyang G, Sommer W, Zhou C (2015b) Updating and validating a new framework for restoring and analyzing latency-variable erp components from single trials with residue iteration decomposition (ride). Psychophysiology 52:839–856.

Ouyang G, Sommer W, Zhou C (2016) Reconstructing erp amplitude effects after compensating for trial-to-trial latency jitter: a solution based on a novel application of residue iteration decomposition. Int J Psychophysiol 109:9–20.

Parsons TD (2015) Virtual reality for enhanced ecological validity and experimental control in the clinical, affective and social neurosciences. Front Hum Neurosci 9:1–19.

Peirce J, Gray JR, Simpson S, MacAskill M, Höchenberger R, Sogo H, Kastman E, Lindeløv JK (2019) PsychoPy2: experiments in behavior made easy. Behav Res Methods 51:195– 203.

Perrin F, García-Larrea L, Mauguière F, Bastuji H (1999) A differential brain response to the subject’s own name persists during sleep. Clin Neurophysiol 110:2153–2164.

Perrin F, Maquet P, Peigneux P, Ruby P, Degueldre C, Balteau E, Del Fiore G, Moonen G, Luxen A, Laureys S (2005) Neural mechanisms involved in the detection of our first name: a combined erps and pet study. Neuropsychologia 43:12–19.

Perrin F, Schnakers C, Schabus M, Degueldre C, Goldman S, Brédart S, Faymonville ME, Lamy M, Moonen G, Luxen A, Maquet P, Laureys S (2006) Brain response to one’s own name in vegetative state, minimally conscious state, and locked-in syndrome. Arch Neurol 63:562–569.

Power AJ, Foxe JJ, Forde EJ, Reilly RB, Lalor EC (2012) At what time is the cocktail party? a late locus of selective attention to natural speech. Eur J Neurosci 35:1497–1503.

Redick TS, Broadway JM, Meier ME, Kuriakose PS, Unsworth N, Kane MJ, Engle RW (2012) Measuring working memory capacity with automated complex span tasks. Eur J Psychol Assess 28:164–171.

Rivenez M, Darwin CJ, Bourgeon L, Guillaume A (2007) Unattended speech processing: effect of vocal-tract length. J Acoust Soc Am 121:EL90–EL95.

Rivenez M, Darwin CJ, Guillaume A (2006) Processing unattended speech. J Acoust Soc Am 119:4027–4040.

Rivenez M, Guillaume A, Bourgeon L, Darwin CJ (2008) Effect of voice characteristics on the attended and unattended processing of two concurrent messages. Eur J Cogn Psychol 20:967–993.

Röer JP, Bell R, Buchner A (2013) Self-relevance increases the irrelevant sound effect: attentional disruption by one’s own name. Eur J Cogn Psychol 25:925–931.

Röer JP, Körner U, Buchner A, Bell R (2017) Attentional capture by taboo words: a functional view of auditory distraction. Emotion 17:740–750.

Schepman A, Rodway P, Pritchard H (2016) Right-lateralized unconscious, but not conscious, processing of affective environmental sounds. Laterality 21:606–632.

Shafiro V, Gygi B (2007) Perceiving the speech of multiple concurrent talkers in a combined divided and selective attention task. J Acoust Soc Am 122:EL229–35.

Shamay-Tsoory SG, Mendelsohn A (2019) Real-life neuroscience: an ecological approach to brain and behavior research. Perspect Psychol Sci 14:841–859.

Shiffrin RM, Schneider W (1977) Controlled and automatic human information processing: ii. perceptual learning, automatic attending and a general theory. Psychol Rev 84:127– 190.

Shinn-Cunningham BG, Ihlefeld A (2004) Selective and divided attention: extracting information from simultaneous sound sources. Proc Int Conf Audit Display, 6-9 July 2004.

Simon JZ (2015) The encoding of auditory objects in auditory cortex: insights from magnetoencephalography. Int J Psychophysiol 95:184–190.

Tacikowski P, Nowicka A (2010) Allocation of attention to self-name and self-face: an erp study. Biol Psychol 84:318–324.

Tamura K, Karube C, Mizuba T, Iramina K (2012) ERP and time frequency analysis of response to subject’s own name. 5th 2012 Biomed Eng Int Conf BMEiCON 2012:12–15.

Tateuchi T, Itoh K, Nakada T (2012) Neural mechanisms underlying the orienting response to subject’s own name: an event-related potential study. Psychophysiology 49:786–791.

Tóth B, Farkas D, Urbán G, Szalárdy O, Orosz G, Hunyadi L, Hajdu B, Kovács A, Szabó BT, Shestopalova LB, Winkler I (2019) Attention and speech-processing related functional brain networks activated in a multi-speaker environment. PLoS One 14:e0212754.

Treisman AM (1960) Contextual cues in selective listening. Q J Exp Psychol 12:242–248.

Tun PA, O’Kane G, Wingfield A (2002) Distraction by competing speech in young and older adult listeners. Psychol Aging 17:453–467.

van der Wulp I (2021) Word segmentation: tp or ocp? a re-analysis of batterink & paller (2017).

Vazquez Alvarez Y, Brewster SA (2010) Designing spatial audio interfaces to support multiple audio streams. ACM Int Conf Proceeding Ser:253–256.

Wood N, Cowan N (1995) The cocktail party phenomenon revisited: how frequent are attention shifts to one’s name in an irrelevant auditory channel? J Exp Psychol Learn Mem Cogn 21:255–260.

Xia J, Nooraei N, Kalluri S, Edwards B (2015) Spatial release of cognitive load measured in a dual-task paradigm in normal-hearing and hearing-impaired listeners. J Acoust Soc Am 137:1888–1898.

Ye H, Fan Z, Chai G, Li G, Wei Z, Hu J, Sheng X, Chen L, Zhu X (2021) Self-related stimuli decoding with auditory and visual modalities using stereo-electroencephalography. Front Neurosci 15:1–15.

Zion Golumbic EM, Ding N, Bickel S, Lakatos P, Schevon CA, McKhann G, Goodman RR, Emerson R, Mehta AD, Simon JZ, Poeppel D, Schroeder CE (2013) Mechanisms underlying selective neuronal tracking of attended speech at a cocktail party . Neuron 77:980–991.

